# Generating functional plasmid origins with OriGen

**DOI:** 10.1101/2025.02.03.636306

**Authors:** Jamie Irvine, Jigyasa Arora, Jonathan N.V. Martinson, Jaymin R. Patel, Sarah I. Hasham, Brady F. Cress, Benjamin E. Rubin

## Abstract

While generative artificial intelligence has shown promise for biological design, no computational system has yet created sequences proven capable of replication. Focusing on plasmids as minimal replicating systems, we develop OriGen, a language model that generates novel plasmid origins of replication while maintaining essential functional elements. We experimentally validate OriGen’s ability to create functional origins that diverge from existing wild types, demonstrating the model’s capacity to capture the complex and often enigmatic mechanisms of biological replication.

## MAIN

In recent years, artificial intelligence (AI) has transformed how we study biological systems, with tools like AlphaFold revealing fundamental insights about molecules and their interactions^1^. While AI has been invaluable for these descriptive tasks, its potential for biological design remains largely untapped. Current bioengineering approaches rely heavily on manual design and laborious trial-and-error, confining our capabilities to a handful of model systems^2,3^. Recent advances in generative modeling, exemplified by groundbreaking achievements in natural language processing^4,5^, suggest a promising path forward. These tools can overcome current limitations by rapidly creating novel sequences for user-defined functions, efficiently exploring vast regions of sequence space that remain otherwise inaccessible. Combined with ever-decreasing synthesis costs, this approach has the potential to dramatically accelerate bioengineering, enabling bespoke solutions for the wide diversity of biological design challenges.

Recent work has shown promising steps toward machine-generated biological sequences. Protein generation models like ProGen^6^ and ProtGPT2^7^ have demonstrated the ability to create novel functional proteins across a range of structural families. The field has also expanded into DNA sequence generation, with large language models like GenSLMs^8^ and Evo^9^ showing increasing capability to generate both coding and non-coding regions in a diversity of settings. But despite growing investment in these DNA foundation models, they are still in their infancy, and have yet to demonstrate mastery of the full spectrum of sequences and functions. Notably, plasmids have received surprisingly little attention from generation efforts, despite being crucial for the transfer and evolution of new genes and essential tools for biological engineering. While there has been some recent work generating plasmid and phage genomes *in silico*, the outputs remain untested in biological systems^10,11^. In particular, no study has experimentally validated AI-generated sequences capable of plasmid replication—the only essential component of a plasmid, and a fundamental aspect of life itself.

Here we address this gap by developing OriGen, a language model specifically trained to generate replicons—the minimal genetic units required for DNA replication. We assemble a comprehensive database of plasmid replicons and their host associations, then use this to train a model that can generate novel replication sequences. OriGen is validated by synthesizing generated origins of replication (*oriVs*) and testing their ability to replicate in bacterial cells. This work represents the first experimentally validated AI-generated biological sequences capable of replication.

OriGen is an autoregressive language model capable of generating host-dependent plasmid replicons, which consist of an *oriV* and Replication initiation (Rep) protein(s) (Fig. 1a). To train this model, we first assembled a comprehensive dataset of *oriVs* and Reps from plasmids, along with their bacterial hosts. This presented a significant challenge, as *oriVs* are poorly characterized and difficult to identify computationally. The only dedicated database of plasmid *oriVs*, DoriC^12^, provides carefully curated sequences, but contains relatively few examples. To expand this dataset, we developed a BLAST-based pipeline to identify putative *oriVs* in two larger plasmid databases: PLSDB^13^ and IMG/PR^14^. By aligning the non-coding region of the genomes from these databases against known *oriVs* from DoriC, this pipeline added 24,568 newly annotated replicons from PLSDB and 21,229 from IMG/PR to the 1,209 examples in DoriC (Fig. 1b). Associated Rep genes were identified using a Rep-specific HMM search^15,16^ and protein database match^17^, creating the largest known collection of plasmid replicon sequences.

**Figure 1:**
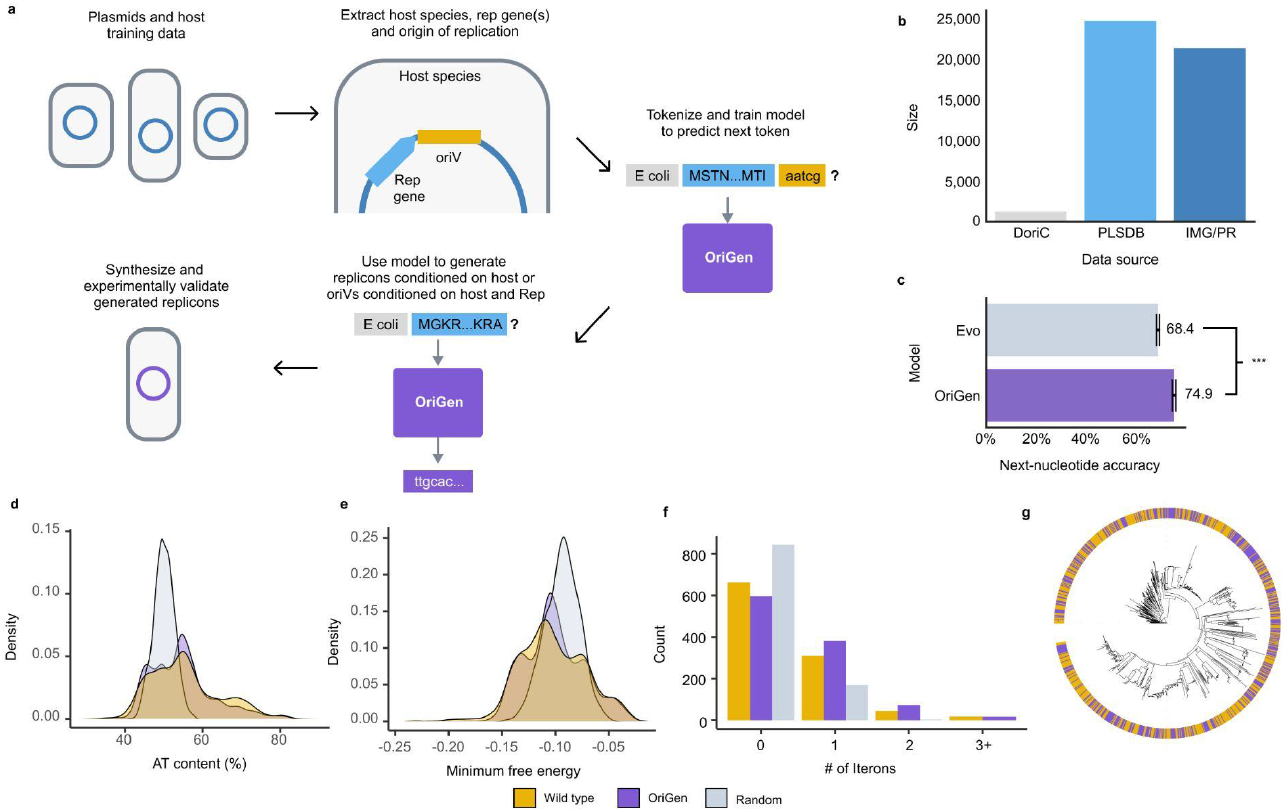
Development and validation of OriGen for plasmid origin generation: **a**, Schematic of the OriGen training, generation, and validation pipeline. **b**, Replicon data extracted from different plasmid datasets. **c**, Next-nucleotide prediction accuracy comparison (two-sided bootstrap test, P < 0.001). **d-g**, Analysis based on 1,000 *oriVs* from unprompted model-generated replicons compared to equal numbers of sampled wild-type *oriVs* and random sequences, showing distributions of: **d**, AT content, **e**, minimum free energy, and **f**, number of iterons, as well as **g**, phylogenetic tree of model-generations vs wild types.

OriGen was trained on this replicon dataset. For each plasmid, we extracted and tokenized three components into a single sequence: the bacterial host species, followed by the amino acid sequence(s) of the Rep protein(s) (when present), followed by the *oriV* nucleotide sequence. To ensure robust evaluation, we combined the DoriC and PLSDB datasets and split them into training and validation splits based on *oriV* sequence similarity, and we reserved the IMG/PR dataset as a held-out test set to evaluate domain-transfer performance across markedly different data sources (Supplemental Figure 1). Once trained, OriGen can generate new replicon sequences from scratch, replicons conditioned for a desired host species compatibility, or *oriVs* conditioned on a given Rep gene.

As an initial validation, we evaluated OriGen’s ability to predict the next nucleotide in *oriV* sequences from our held-out IMG/PR test set. To ensure no near-duplicates of our training set were in our test set, we excluded any sequences with greater than 95% similarity (BLAST pident > 95%, qcovs > 95%) to those in our training data. We compared OriGen’s performance to Evo, which has recently emerged as a prominent genomic language model. Unlike OriGen, Evo was trained on IMG/PR itself, making direct comparison difficult. Moreover, Evo’s train/test splits were unavailable at the time of analysis, so a substantial portion of our evaluation set is likely part of Evo’s training data. Despite this considerable disadvantage, OriGen achieved significantly higher accuracy in predicting *oriV* sequences (75% vs 68%, Fig. 1c; bootstrap 95% CIs: [74.2-75.5] vs [67.8-69.0]). This performance gap suggests that specialized models can outperform general foundation models for specific biological tasks like replicon generation.

Beyond next-nucleotide prediction, we also analyzed the characteristics of unprompted model-generated replicon sequences to assess their biological plausibility. We generated 1,000 sequences with no prompt (i.e. no specified species or Rep) and compared their *oriV* regions to those from wild-type plasmids and random sequences. While natural *oriV* sequences are diverse and not fully characterized, they often contain recognizable features such as AT-rich regions, low minimum free energy, and repeated sequences called iterons, which facilitate DNA replication initiation. The distribution of these motifs in our generated sequences closely matched that of wild-type *oriVs*, while random sequences showed markedly different patterns (Fig. 1d-f). Additionally, we built a phylogenetic tree of the generated and wild-type origins together, revealing that model-generated *oriVs* are distributed throughout all major branches alongside wild-type *oriVs* (Fig. 1g). This indicates that the model generates diverse sequences rather than variations of a single template.

To validate that OriGen generates functional replication sequences, we tested their performance in bacterial cells (Fig. 2a). To test generated *oriVs*, we synthesized and assembled them into vectors (R6K) that are dependent on a host gene (*pir*) for replication. The resulting constructs were transformed in parallel into *pir+* (EC100D) and *pir-* (NEB 10-beta) strains. The *pir+* strain served as an assembly control, as these cells would maintain the plasmid via the *R6K* origin regardless of the synthetic origin’s functionality. In contrast, growth in the *pir-* strain should only occur if our generated origin functioned. We confirmed the identity and integrity of constructs through full plasmid sequencing, allowing us to confirm the validity of the experiment and detect any mutations that arose during synthesis, assembly, or replication, which may indicate a “near miss” sequence that is one nucleotide away from being functional.

**Figure 2:**
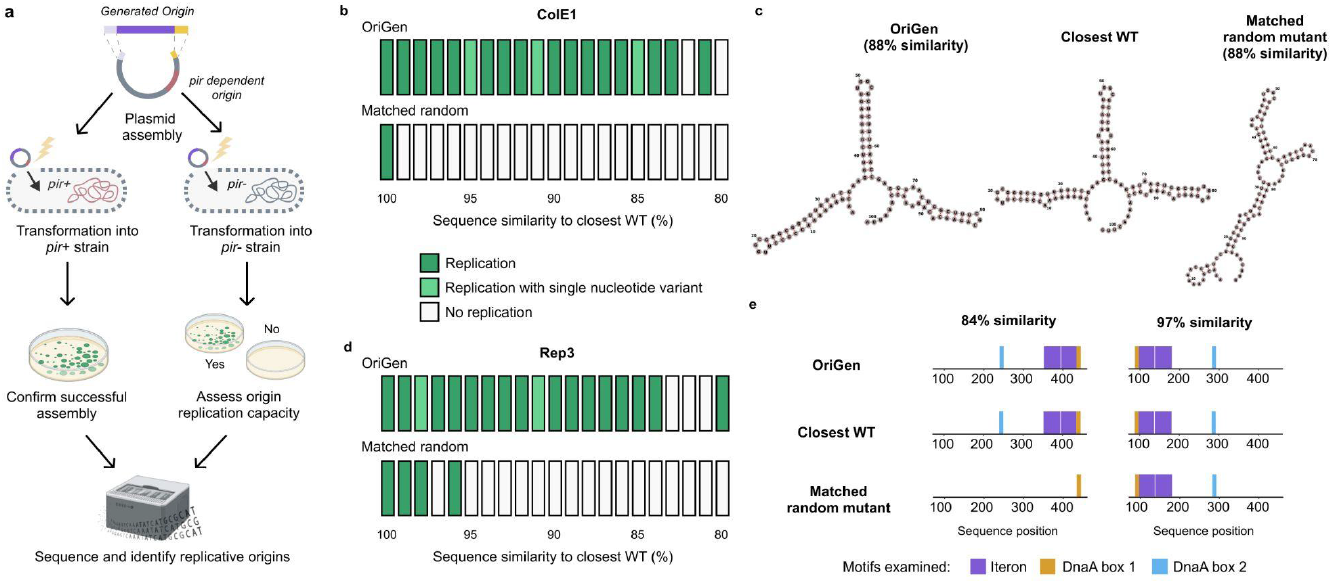
Functional validation of OriGen-generated origins: **a**, Experimental workflow for validating generated origins. **b**, Assessment of ColE1-type origin function in *E. coli*. Matched random sequences contain the same number of mutations relative to wild-type as their corresponding OriGen-generated sequences. Lighter green indicates a single nucleotide polymorphism or single nucleotide indel was detected after sequencing. **c**, RNAI secondary structure comparison of one OriGen-generated ColE1 example vs closest WT vs matched random mutant control. **d**, Assessment of uncharacterized Rep3-type origin function in *E. coli*. **e**, Distribution of known motifs in examples of Rep3 OriGen-generated origins vs closest WT vs matched random mutant. In both examples, the model-generated origins led to replication while the random mutants did not.

For our first experimental test, we focused on ColE1-type replicons, which are among the most thoroughly studied plasmid replication systems^18^. ColE1 origins include two overlapping antisense RNAs^19,20^, and typically function without a plasmid-encoded Rep protein. We prompted OriGen with an *E. coli* species token, no Rep protein sequence, and a 20-base seed sequence from a randomly selected ColE1 origin in our validation set. From 100,000 generated origins, we sampled candidates with varying levels of similarity to known origins through sequence alignment, spanning from 99% down to 80% similarity to their closest match in the training data. These generated sequences enabled plasmid replication across multiple similarity levels, with successful replication observed at similarities as low as 81% (Fig. 2b). In several cases (at 95%, 91%, and 85% similarity), sequencing revealed single base pair changes in the replicated plasmids. To test whether our success reflected genuine functional design rather than simple robustness to mutation, we created control sequences by introducing matched numbers of random mutations into the wild-type origins most closely related to the generated sequence. None of these randomly mutated controls enabled replication, suggesting that OriGen learned to preserve essential functional elements even in a replication system highly sensitive to mutations.

To understand how OriGen preserved functionality while generating novel sequences, we examined trios of related sequences: model-generated origins, their nearest wild-type matches, and corresponding randomly mutated versions. We analyzed RNAI secondary structures, a well-characterized regulatory element^21,22^ that forms a cloverleaf structure that inhibits ColE1 replication, providing copy number control^23^ (Fig. 2c). Generated sequences generally maintained their characteristic secondary structure despite multiple nucleotide differences from their wild-type counterparts. In contrast, in the matching randomly mutated sequences, similar numbers of differences typically disrupted the cloverleaf structure (Supplemental Figure 2). Despite training only on DNA sequences, OriGen learned to navigate the inherent structural sensitivity of origins by strategically introducing variations that preserve critical functional features.

To evaluate OriGen’s performance beyond well-characterized replicons, we also tested its ability to generate origins for a naturally occurring but previously uncharacterized plasmid. We identified an *E. coli* plasmid with no documented laboratory use and prompted OriGen with its Rep protein (Rep3) sequence to generate compatible *oriVs*. Following the same experimental pipeline as with ColE1, we cloned these generated origins alongside the Rep gene into our R6K testing system. We again observed successful replication across multiple similarity levels, with generated sequences functioning down to 80% similarity to known origins (Fig. 2d).

Analysis of Rep3 sequence trios revealed patterns in the conservation of known functional motifs (Fig. 2e). In most cases, matched random mutations disrupted essential elements like iterons and DnaA boxes, while OriGen preserved them, directly explaining the differences in functionality. More intriguingly, in another case, both the generated and randomly mutated sequences maintained all known motifs, yet only the model-generated sequence enabled replication (97% similarity sequences, Fig. 2e, Supplemental Figure 3). This suggests that beyond the currently characterized motifs, there are additional sequence features essential for replication that our model learned to preserve. OriGen’s ability to maintain these uncharacterized but crucial elements, even while generating substantially different sequences, demonstrates the potential for AI systems to capture complex biological design constraints that extend beyond our current understanding.

Interestingly, while the generated ColE1 sequences differed from associated wild types in positions scattered throughout the origin, the variations in the Rep3 generated origins were more concentrated in the terminal 60 bp at the 3’ end of the sequence (Supplemental Figure 4). The model seems to be signaling a non-essential region of the Rep3 origin, suggesting its ability to be used for origin boundary detection. To validate this, we removed the terminal 60 bp from a wild-type Rep3 origin and found that the plasmid replicated (Extended Data S4).

In this work, we have developed and experimentally validated a language model capable of generating functional origins of replication. By assembling a comprehensive dataset of plasmid replicons and their bacterial hosts, we trained a model that learned to preserve the complex requirements for plasmid replication while exploring variations in sequence. Our experimental validation in *E. coli* across two fundamentally different replication mechanisms demonstrates this capability and suggests a generalizable understanding of replication.

While this work demonstrates promising results, several limitations should be noted. Our training dataset, while substantially larger than previous collections, captures only a subset of replicon diversity. This constraint stems from our computational pipeline, which necessarily restricts us to sequences similar to documented origins in DoriC, likely missing undiscovered replication mechanisms present in nature. Our experimental validation centered on *E. coli* as a model system. Though our architecture incorporates host information, testing host-specificity across diverse microbes remains as future work. Additionally, while OriGen successfully generates functional sequences, the black-box nature of neural networks limits our ability to extract explicit rules about replicon function. Finally, like all machine learning systems, OriGen generates sequences derived from patterns in its training data, producing novelty from combinations and variations of natural sequence elements, rather than inventing entirely new mechanisms.

Despite these caveats, this work represents the first example of AI-generated sequences capable of biological replication. As such, it is important to consider potential safety implications. While the origins generated in this study differ measurably from naturally occurring sequences, they do share known conserved features and are unlikely to present novel biosafety concerns beyond standard laboratory plasmid work. Additionally, we followed established safety protocols to prevent environmental release of recombinant DNA. However, as AI systems become increasingly capable of generating more diverse biological sequences, careful consideration of ethical, legal, and social implications will be essential. This work provides an opportunity to begin discussions about appropriate safety frameworks while these technologies are still in their early stages.

The ability to generate functional replication computationally enables new opportunities for biological design. This work opens the path to the creation of origins functional in diverse bacterial hosts, which is of particular interest for the broad majority of bacteria that lack robust genetic tools. Extending OriGen’s capabilities to whole plasmid generation could facilitate targeted microbiome editing across clinical, industrial, and agricultural applications. This would allow researchers to quickly move from sequencing the community to design, synthesis, and deployment of host-compatible plasmids functionally tailored to address undesirable microbiome characteristics. While significant technical hurdles remain to achieve this vision, OriGen represents a step toward programming biological replication for human-defined purposes.

## METHODS

### Training data generation

To expand the plasmid replicon database, the plasmid genomes in the PLSDB^13^ (version 2023_11_23_v2) and IMG/PR^14^ were annotated using Prokka^24^ (version 1.14.6) and Prodigal^25^ (version 2.6.3) software. The non-coding regions were defined from these annotations using BEDTools suite software toolkit^26^ (version 2.31.1). All the protein sequences were annotated against the PFAM database^15^ (version 33.1) using hmmsearch (version 3.4, http://hmmer.org/). The Rep proteins were annotated by their PFAM domains. These annotations were supplemented with custom Rep HMMs generated by the plaSquid software^16^ and BLASTn against the MOB-suite Replication protein database^17^.

The non-coding regions were annotated by BLASTn against the DoriC database^12^ (version 12.0) and filtered for 80% identity and 80% coverage. To prevent misclassification, BLASTn-selected regions were further filtered based on their proximity to Rep proteins.

Host assignments were inferred from isolation source and CRISPR spacer matching data within PLSDB and IMG/PR^13,14^.

### Model training

Training and validation splits were created by first clustering *oriV* sequences from the combined DoriC and PLSDB datasets using CD-HIT (v4.8.1) with a 90% sequence identity threshold. The resulting clusters were then randomly split 75/25 for training and validation, with the validation set used for hyperparameter tuning and early stopping. The IMG/PR dataset was reserved as a separate held-out test set, after excluding any *oriVs* with greater than 95% similarity (BLAST pident > 95%, qcovs > 95%) to any *oriV* in our training split.

Input sequences were tokenized as discrete units: bacterial host species names as single tokens, amino acids for Rep proteins, and individual nucleotides for *oriV* sequences (using lowercase to avoid collisions with amino acid letters). Special tokens handled cases of unknown or unannotated species and separation of multiple Rep proteins when present. We chose amino acid-level tokenization for Rep proteins primarily to reduce sequence lengths within our maximum context window of 1,500 tokens. The complete vocabulary contained 145 tokens.

OriGen uses a decoder-only transformer architecture with 12 layers and 12 attention heads per layer (embedding dimension 768), totaling approximately 86M parameters. The model was trained using AdamW optimizer with standard dropout rates (0.1) and weight decay (0.1). Training used mixed precision and ran on a single NVIDIA RTX A5000 GPU, with early stopping based on validation loss.

### Model evaluation

To enable direct comparison between OriGen and Evo (8k base model), which uses single-nucleotide tokenization on contiguous DNA sequences, we adapted our test sequences accordingly. For each plasmid in the IMG/PR dataset, we extracted the contiguous DNA region containing both the Rep gene and its adjacent *oriV*. Since our evaluation focuses on *oriV* prediction conditioned on Rep sequence, we took the reverse complement of sequences where the Rep gene followed the *oriV*, ensuring Rep always preceded *oriV* while maintaining natural sequence contexts. Test sequences were filtered to exclude any with greater than 95% similarity to sequences in our training data (BLAST parameters: pident > 95%, qcovs > 95%). Since Evo’s train/test splits are not currently published, the majority of sequences in this evaluation were likely part of Evo’s training data. Prediction accuracy was calculated as the percentage of correctly predicted nucleotides across all test sequences, with 95% confidence intervals estimated through bootstrapping (n=1000 resamples).

### Model generations

For the analysis of unprompted replicon generation (Fig. 1d-g), sequences were sampled using nucleus sampling (top-p=0.95) with temperature 1.0. For experimental validation (Fig. 2), both replicon types were conditioned with a starting *E. coli* token. The ColE1 examples were further conditioned with the first 20 nucleotides from a ColE1 *oriV* from the validation split (with no amino acid tokens). The Rep3 sequences were conditioned with the amino acid sequence from the Rep gene identified in the chosen plasmid. These generations used top-k sampling (k=4) with temperature 1.0. In both cases, generation proceeded until the model produced a stop token or reached the maximum sequence length.

### In-silico validation

To compare OriGen-generated sequences with wild-type and random sequences (Fig. 1d-g), we generated 1,000 unprompted sequences from OriGen and extracted their *oriVs*. We compared these with 1,000 wild-type *oriVs* randomly sampled from the training data. As a baseline, 1,000 random nucleotide sequences were generated, each of length 357, the median *oriV* length in the training data.

The occurrence of known conserved motifs in *oriVs* was searched using FIMO software from the MEME suite^27^. These motifs were identified via a literature search of known motifs in the *oriVs* such as DnaA boxes and iterons. The annotated motifs were filtered based on a log-likelihood score greater than 1.0 and the presence of all motifs within the region. AT content was calculated for each sequence using an in-house Python script. Minimum free energy (MFE) was calculated on a sliding window of 100 base pairs using RNAfold (version 2.5.1) from the ViennaRNA package^28^. The sliding window was created using Seqkit^29^ (version 2.9.0). Mean MFE scores were calculated and plotted for each sequence. All sequences were aligned using MAFFT^30^ (version 7.505), and a phylogenetic tree was constructed using FastTree with the GTR algorithm^31^. The tree was visualized in iTOL^32^.

### Sequence similarity and matched random mutants

To experimentally test generated sequences with varying levels of novelty, we quantified sequence similarity using global alignment (Needleman–Wunsch algorithm, implemented via EMBOSS needle with gap_open=10.0 and gap_extend=0.5). For each generated sequence, we first identified candidate wild-type matches by BLAST search against our training data, then performed global alignment with the top BLAST hit to calculate a final similarity score. (Supplemental Table 1)

To create matched random mutants as controls, we analyzed these alignments to determine the number of insertions, deletions, and point mutations between each generated sequence and its closest wild-type match. We then created control sequences by applying the same number of each mutation type (e.g., insertion, deletion, SNP) at random positions in the wild-type sequence. Due to the interaction between mutations, global alignment of these random controls back to their wild-type sequences sometimes yielded different optimal alignments, resulting in similarity scores that could be slightly higher or lower than their model-generated counterparts.

### Experimental sequence analysis

RNAI and RNAII regions of the tested ColE1 replicons were annotated using HMM profiles from a previous publication^33^. The longest aligned RNAI region was extracted using the samtools faidx command^34^. RNAI secondary structures were predicted for the extracted RNAI sequences using the RNAfold WebServer^28^. Conserved motifs in experimentally tested Rep3-containing *oriVs* were identified using the FIMO software from the MEME suite^27^ as described above.

### Bacterial strains and culture conditions

Two bacterial strains were used in this study. Transformax EC100D *pir*+ (Epicenter, Cat. No. 75927-934) has the genotype *F*^−^ *mcrA* Δ*(mrr-hsdRMS-mcrBC) φ80dlacZ*Δ*M15* Δ*lacX74 recA1 endA1 araD139* Δ*(ara, leu)7697 galU galK λ*^−^ *rpsL (Str^R^) nupG pir*^+^*(DHFR)*. This strain was selected for its ability to maintain and propagate plasmids requiring the *pir* gene for replication such as suicide plasmids with *R6K* origins. NEB 10-Beta (New England Biolabs, Cat. No. C3020K) has the genotype Δ*(ara-leu)7697 araD139 fhuA* Δ*lacX74 galK16 galE15 e14*^−^ *φ80dlacZ*Δ*M15 recA1 relA1 endA1 nupG rpsL (Str^R^) rph spoT1* Δ*(mrr-hsdRMS-mcrBC)* and was chosen for its high transformation efficiency and lack of *pir*. Bacterial cultures were grown in lysogeny broth (LB; RPI, Cat. No. L24060-5000.0) supplemented with carbenicillin (50 µg/mL; Sigma-Aldrich, Cat. No. C1389) at 37°C with shaking at 200 rpm. For solid media, lysogeny broth containing 2% agar (BD, Cat. No. 214010) was used.

### Plasmid construction

The R6K suicide vector backbone used in this study was adapted from a previously developed part plasmid^35^. All PCRs were performed using Q5® High-Fidelity 2X Master Mix (New England Biolabs, Cat. No. M0492L) following the manufacturer’s instructions.

To insert the ColE1 origins, BsaI Golden Gate recognition sites were added to an in-house R6K vector via PCR amplification using oligonucleotides oJNVM0070 and oJNVM0073 (Extended Data S1 and S2). PCR amplicons were gel purified with the Zymoclean Gel DNA Recovery Kit (Zymo Research, Cat. No. D4002). ColE1 origins were synthesized as gene fragments (Twist Biosciences) containing BsaI sites and complementary overhangs (Extended Data S3).

For the construction of the rep-associated origin vector, Gibson assembly was used to clone the Rep gene under the control of a weak promoter (BBa_J23114) and a ribosomal binding site designed with the “Control Translation” function on denovodna.com at a target translation efficiency rate of 50,000 a.u.^36^. Additionally, a SapI Golden Gate dropout cassette was inserted immediately downstream of the Rep gene, providing a site for the insertion of replication origins (Extended Data S2). DNA fragments encoding the promoter, RBS, Rep gene, and dropout cassette were synthesized as a single gene fragment (Twist Biosciences) with overhangs homologous to the R6K vector.

Initial experiments indicated that wild-type origins of replication functioned only in the presence of a P5R mutation in the *rep3* gene. This mutation was introduced into the Rep dropout cassette vector using site-directed mutagenesis.

Golden Gate assemblies were performed using type IIS restriction enzymes BsaI-HF®v2 (New England Biolabs, Cat. No. R3733S) for ColE1 origins and SapI (New England Biolabs, Cat. No. R0569S) for rep-associated origins. Each reaction was performed with a final concentration of 4 nM vector and insert.

Gibson assemblies were performed using NEBuilder® HiFi DNA Assembly Master Mix (New England Biolabs, Cat. No. E2621L) following the manufacturer’s instructions.

### Transformation and screening of plasmids

Assemblies were transformed into either NEB 10-Beta or EC100D *pir*+ cells using electroporation with 0.1 cm Pulser/MicroPulser Electroporation Cuvettes (Bio-Rad, Cat. No. 1652083). Following electroporation, cells were recovered in SOC media (Invitrogen, Cat. No. 15544034) at 37°C for 30 minutes under static conditions. Recovered cells were then centrifuged at 3000 × g for 3 minutes, the supernatant was removed, and the entire cell pellet was streaked onto selective media plates.

Well-separated transformant colonies were picked and inoculated into 5 mL of lysogeny broth containing carbenicillin and incubated overnight at 37ºC with shaking. Freezer stocks were prepared in LB containing 10% glycerol (Sigma-Aldrich, Cat. No. G33-1).

Plasmid DNA was extracted using the QIAprep Spin Miniprep Kit (Qiagen, Cat. No. 27106) and eluted with 30 µL molecular-grade water. DNA concentrations were quantified using a NanoDrop 2000 spectrophotometer, and plasmids were submitted for Oxford Nanopore plasmid sequencing (Plasmidsaurus). Assembled plasmid DNA sequences were aligned to reference sequences using SnapGene (version 8.0.1; Extended Data S4).

## DATA AVAILABILITY

Source code for model training, inference, and evaluation is available on GitHub (https://github.com/j-irvine/origen). The repository includes scripts for sequence generation, evaluation metrics, alignment utilities, and example notebooks demonstrating usage and the sequence similarity analysis pipeline. Pre-trained models and dataset files are publicly hosted on HuggingFace (https://huggingface.co/jirvine). Sequences of experimentally validated origins are provided in the supplementary information.

## ACKNOWLEDGEMENTS

We thank Honglue Shi for advice on RNA structural analysis and Naoko Tanese for suggestions about biological interpretations. We acknowledge support from the Berkeley Initiative for Optimized Microbiome Editing (BIOME), particularly from Jennifer Doudna, Jill Banfield, Brad Ringeisen, Audrey Glynn, and Rachel K. Evans. We also thank undergraduates Christine Corry, Danh Tran, and Emily Qi for their assistance in developing OriGen.

We’d also like to thank our funders. The Audacious Project: This work was supported in part by Lyda Hill Philanthropies, Acton Family Giving, the Valhalla Foundation, Hastings/Quillin Fund - an advised fund of the Silicon Valley Community Foundation, the CH Foundation, Laura and Gary Lauder and Family, the Sea Grape Foundation, the Emerson Collective, Mike Schroepfer and Erin Hoffman Family Fund - an advised fund of Silicon Valley Community Foundation, the Anne Wojcicki Foundation through The Audacious Project at the Innovative Genomics Institute; Leona M. and Harry B. Helmsley Charitable Trust: This work was funded in part by grant [G-2302-06692] from The Leona M. and Harry B. Helmsley Charitable Trust; JBEI: The work conducted, in part, through Joint BioEnergy Institute was supported by the U.S. Department of Energy, Office of Science, Biological and Environmental Research Program, through contract DE-AC02-05CH11231 between Lawrence Berkeley National Laboratory and the U.S. Department of Energy. The United States Government retains and the publisher, by accepting the article for publication, acknowledges that the United States Government retains a nonexclusive, paid-up, irrevocable, worldwide license to publish or reproduce the published form of this manuscript, or allow others to do so, for United States Government purposes. Any subjective views or opinions that might be expressed in the paper do not necessarily represent the views of the U.S. Department of Energy or the United States Government; Shurl and Kay Curci Foundation: This work was also supported by a Research Award from the Shurl and Kay Curci Foundation (https://curcifoundation.org) to the Innovative Genomics Institute Genomic Tool Discovery Program at UC Berkeley, awarded to B.E.R. Funding for open access charge: [JBEI: DE-AC02-05CH11231];; m-CAFEs: This material is funded by m-CAFEs Microbial Community Analysis & Functional Evaluation in Soils, a Science Focus Area led by Lawrence Berkeley National Laboratory is based upon work supported by the U.S. Department of Energy, Office of Science, Office of Biological & Environmental Research under contract number [DE-AC02-05CH11231]; Innovative Genomics Institute: This work was supported in part by the Innovative Genomics Institute; Corundum Convergence Institute: This work was supported by the Corundum Convergence Institute (CCI).

## REFERENCES

1. Jumper, J. et al. Highly accurate protein structure prediction with AlphaFold. Nature 596, 583–589 (2021).

2. Kim, D. et al. Directed evolution and identification of control regions of ColE1 plasmid replication origins using only nucleotide deletions. J. Mol. Biol. 351, 763–775 (2005).

3. Bok, J. W. & Keller, N. P. Fast and easy method for construction of plasmid vectors using modified quick-change mutagenesis. Methods Mol. Biol. 944, 163–174 (2012).

4. Abramson, J. et al. Accurate structure prediction of biomolecular interactions with AlphaFold 3. Nature 630, 493–500 (2024).

5. Lin, Z. et al. Evolutionary-scale prediction of atomic-level protein structure with a language model. Science 379, 1123–1130 (2023).

6. Madani, A. et al. Large language models generate functional protein sequences across diverse families. Nat. Biotechnol. 41, 1099–1106 (2023).

7. Ferruz, N., Schmidt, S. & Höcker, B. ProtGPT2 is a deep unsupervised language model for protein design. Nat. Commun. 13, 4348 (2022).

8. Zvyagin, M. et al. GenSLMs: Genome-scale language models reveal SARS-CoV-2 evolutionary dynamics. Int. J. High Perform. Comput. Appl. 37, 683–705 (2023).

9. Nguyen, E. et al. Sequence modeling and design from molecular to genome scale with Evo. (2024) doi:10.1101/2024.02.27.582234.

10. Shao, B. PlasmidGPT: a generative framework for plasmid design and annotation. bioRxiv 2024.09.30.615762 (2024) doi:10.1101/2024.09.30.615762.

11. Shao, B. & Yan, J. A long-context language model for deciphering and generating bacteriophage genomes. Nat. Commun. 15, 9392 (2024).

12. Dong, M.-J., Luo, H. & Gao, F. DoriC 12.0: an updated database of replication origins in both complete and draft prokaryotic genomes. Nucleic Acids Res. 51, D117–D120 (2023).

13. Schmartz, G. P. et al. PLSDB: advancing a comprehensive database of bacterial plasmids. Nucleic Acids Res. 50, D273–D278 (2022).

14. Camargo, A. P. et al. IMG/PR: a database of plasmids from genomes and metagenomes with rich annotations and metadata. Nucleic Acids Res. 52, D164–D173 (2024).

15. Mistry, J. et al. Pfam: The protein families database in 2021. Nucleic Acids Res. 49, D412–D419 (2021).

16. Giménez, M., Ferrés, I. & Iraola, G. Improved detection and classification of plasmids from circularized and fragmented assemblies. bioRxiv 2022.08.04.502827 (2022) doi:10.1101/2022.08.04.502827.

17. Robertson, J., Bessonov, K., Schonfeld, J. & Nash, J. H. E. Universal whole-sequence-based plasmid typing and its utility to prediction of host range and epidemiological surveillance. Microb. Genom. 6, (2020).

18. Hershfield, V., Boyer, H. W., Yanofsky, C., Lovett, M. A. & Helinski, D. R. Plasmid ColEl as a molecular vehicle for cloning and amplification of DNA. Proc. Natl. Acad. Sci. U. S. A. 71, 3455–3459 (1974).

19. Camps, M. Modulation of ColE1-like plasmid replication for recombinant gene expression. Recent Pat. DNA Gene Seq. 4, 58–73 (2010).

20. Masukata, H. & Tomizawa, J. Control of primer formation for ColE1 plasmid replication: conformational change of the primer transcript. Cell 44, 125–136 (1986).

21. Lacatena, R. M. & Cesareni, G. Base pairing of RNA I with its complementary sequence in the primer precursor inhibits ColE1 replication. Nature 294, 623–626 (1981).

22. Chaillou, S. et al. Directed evolution of colE1 plasmid replication compatibility: a fast tractable tunable model for investigating biological orthogonality. Nucleic Acids Res. 50, 9568–9579 (2022).

23. Cesareni, G., Helmer-Citterich, M. & Castagnoli, L. Control of ColE1 plasmid replication by antisense RNA. Trends Genet. 7, 230–235 (1991).

24. Seemann, T. Prokka: rapid prokaryotic genome annotation. Bioinformatics 30, 2068–2069 (2014).

25. Hyatt, D. et al. Prodigal: prokaryotic gene recognition and translation initiation site identification. BMC Bioinformatics 11, 119 (2010).

26. Quinlan, A. R. & Hall, I. M. BEDTools: a flexible suite of utilities for comparing genomic features. Bioinformatics 26, 841–842 (2010).

27. Grant, C. E., Bailey, T. L. & Noble, W. S. FIMO: scanning for occurrences of a given motif. Bioinformatics 27, 1017–1018 (2011).

28. Lorenz, R. et al. ViennaRNA Package 2.0. Algorithms Mol. Biol. 6, 26 (2011).

29. Shen, W., Le, S., Li, Y. & Hu, F. SeqKit: A cross-platform and ultrafast toolkit for FASTA/Q file manipulation. PLoS One 11, e0163962 (2016).

30. Katoh, K., Misawa, K.Kuma, K.-I. & Miyata, T. MAFFT: a novel method for rapid multiple sequence alignment based on fast Fourier transform. Nucleic Acids Res. 30, 3059–3066 (2002).

31. Price, M. N., Dehal, P. S. & Arkin, A. P. FastTree 2--approximately maximum-likelihood trees for large alignments. PLoS One 5, e9490 (2010).

32. Letunic, I. & Bork, P. Interactive Tree of Life (iTOL) v6: recent updates to the phylogenetic tree display and annotation tool. Nucleic Acids Res. 52, W78–W82 (2024).

33. Ares-Arroyo, M., Rocha, E. P. C. & Gonzalez-Zorn, B. Evolution of ColE1-like plasmids across γ-Proteobacteria: From bacteriocin production to antimicrobial resistance. PLoS Genet. 17, e1009919 (2021).

34. Li, H. et al. The Sequence Alignment/Map format and SAMtools. Bioinformatics 25, 2078–2079 (2009).

35. Liu, H. et al. Magic pools: Parallel assessment of transposon delivery vectors in bacteria. mSystems 3, (2018).

36. Salis, H. M., Mirsky, E. A. & Voigt, C. A. Automated design of synthetic ribosome binding sites to control protein expression. Nat. Biotechnol. 27, 946–950 (2009).

