## Supplemental for "Generating functional plasmid origins with OriGen"

Supplemental Figures

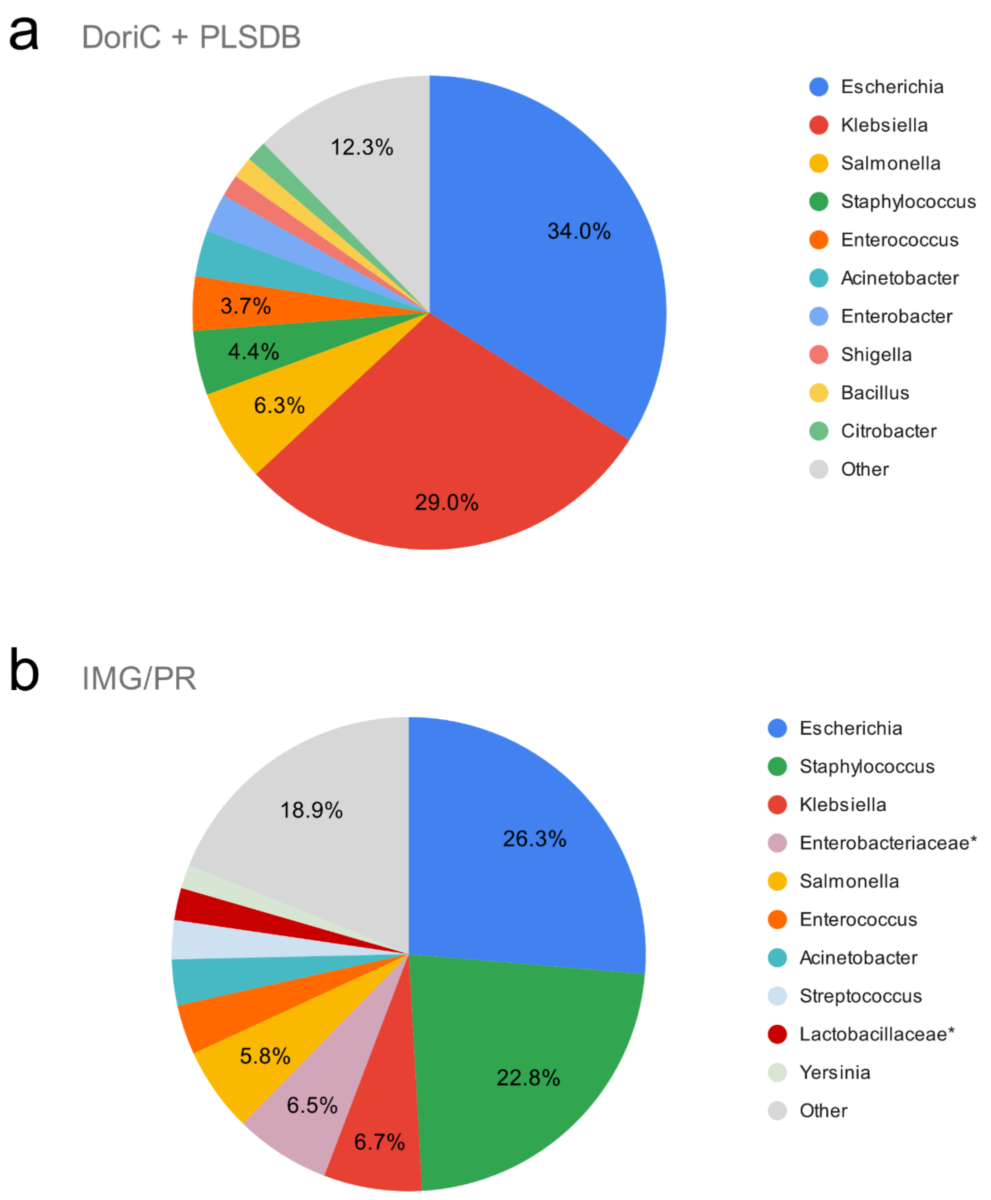


**Supplemental Figure 1: Distribution of host genera in data sets. a,** genera present in DoriC + PLSDB and **b,** IMG/PR. Asterisks indicate where host taxonomy is only known at a higher taxonomic rank.

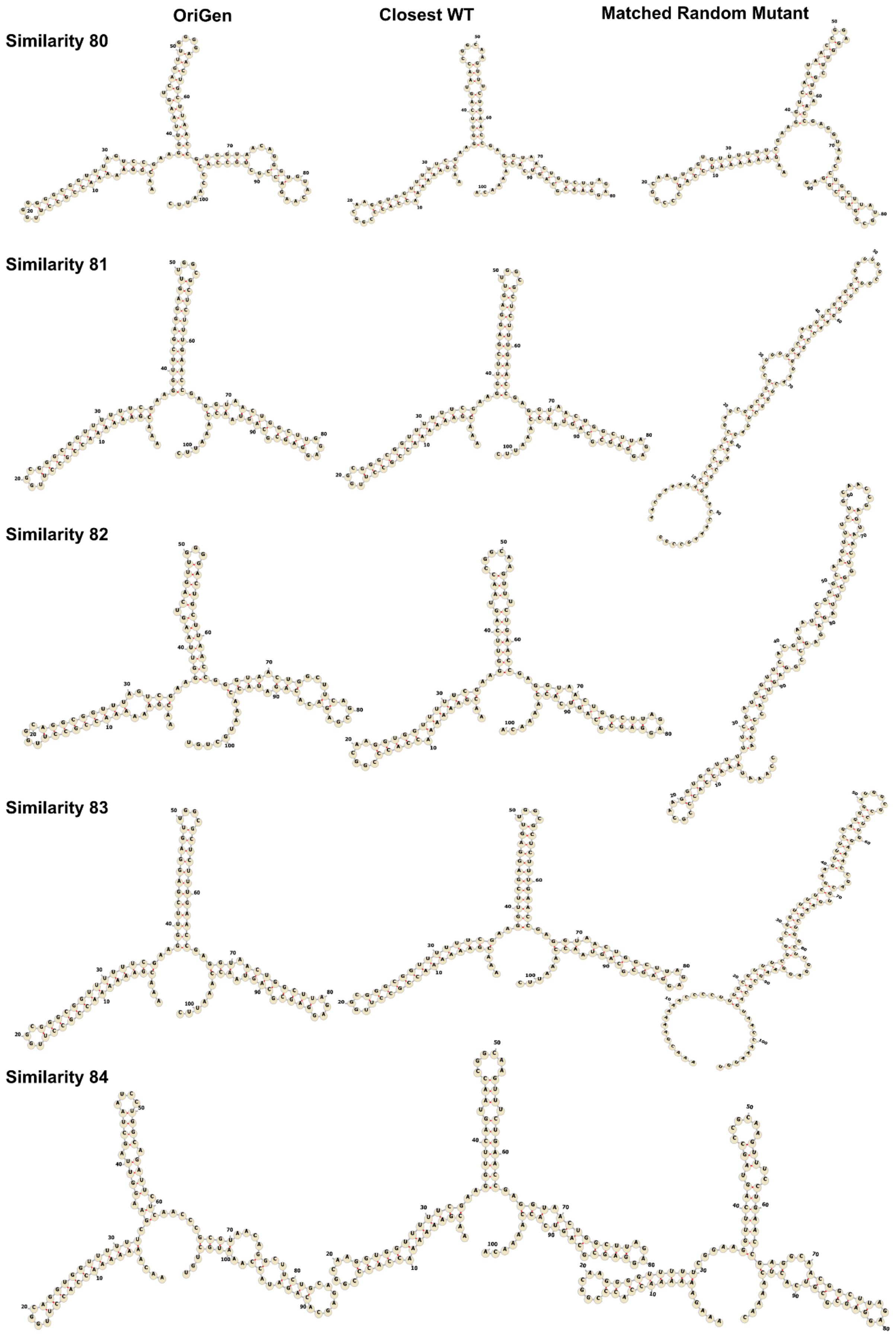

OriGen
Closest WT
Matched Random Mutant
Similarity 80
Similarity 81
Similarity 82
Similarity 83
Similarity 84

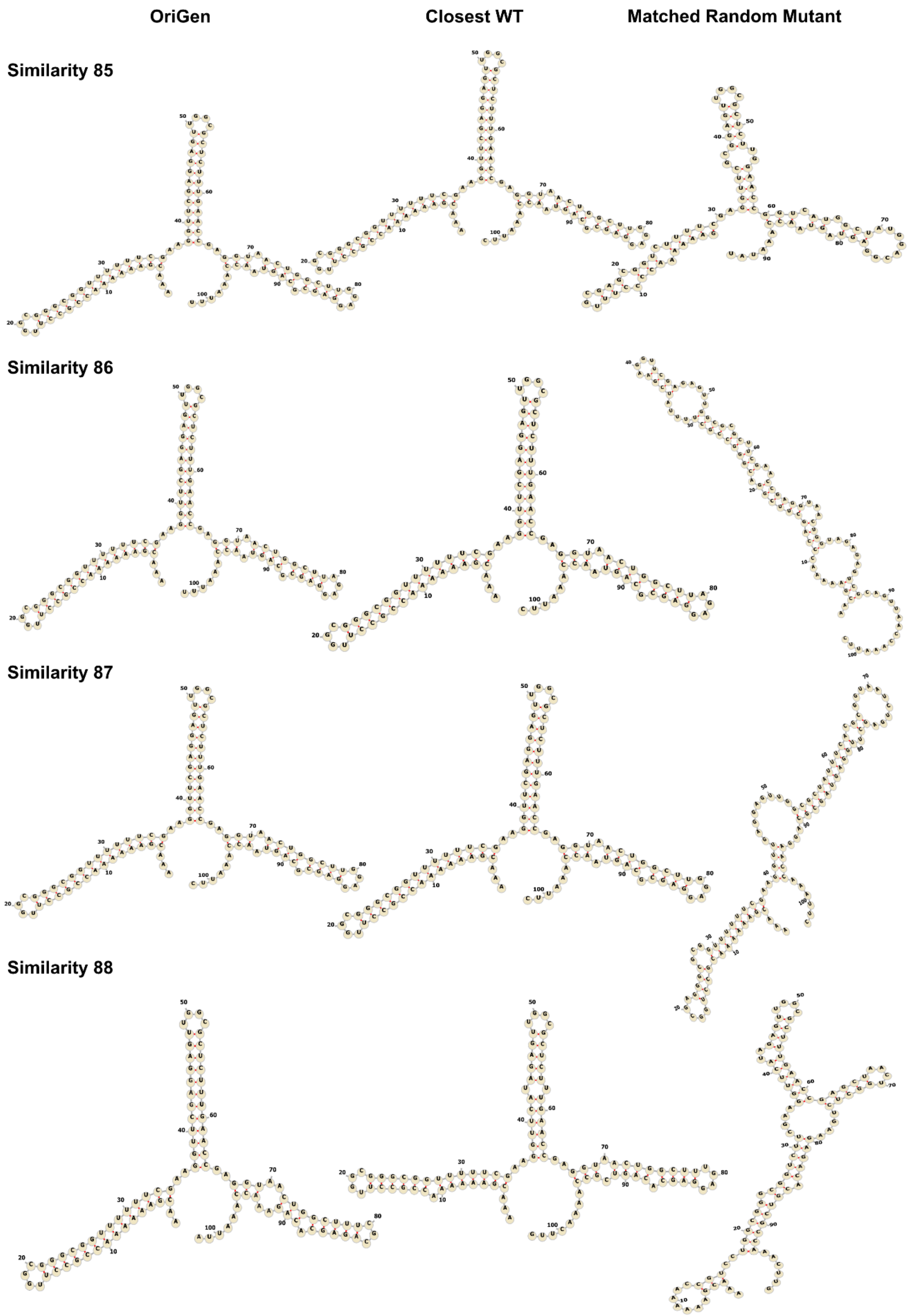

OriGen
Closest WT
Matched Random Mutant
Similarity 85
Similarity 86
Similarity 87
Similarity 88

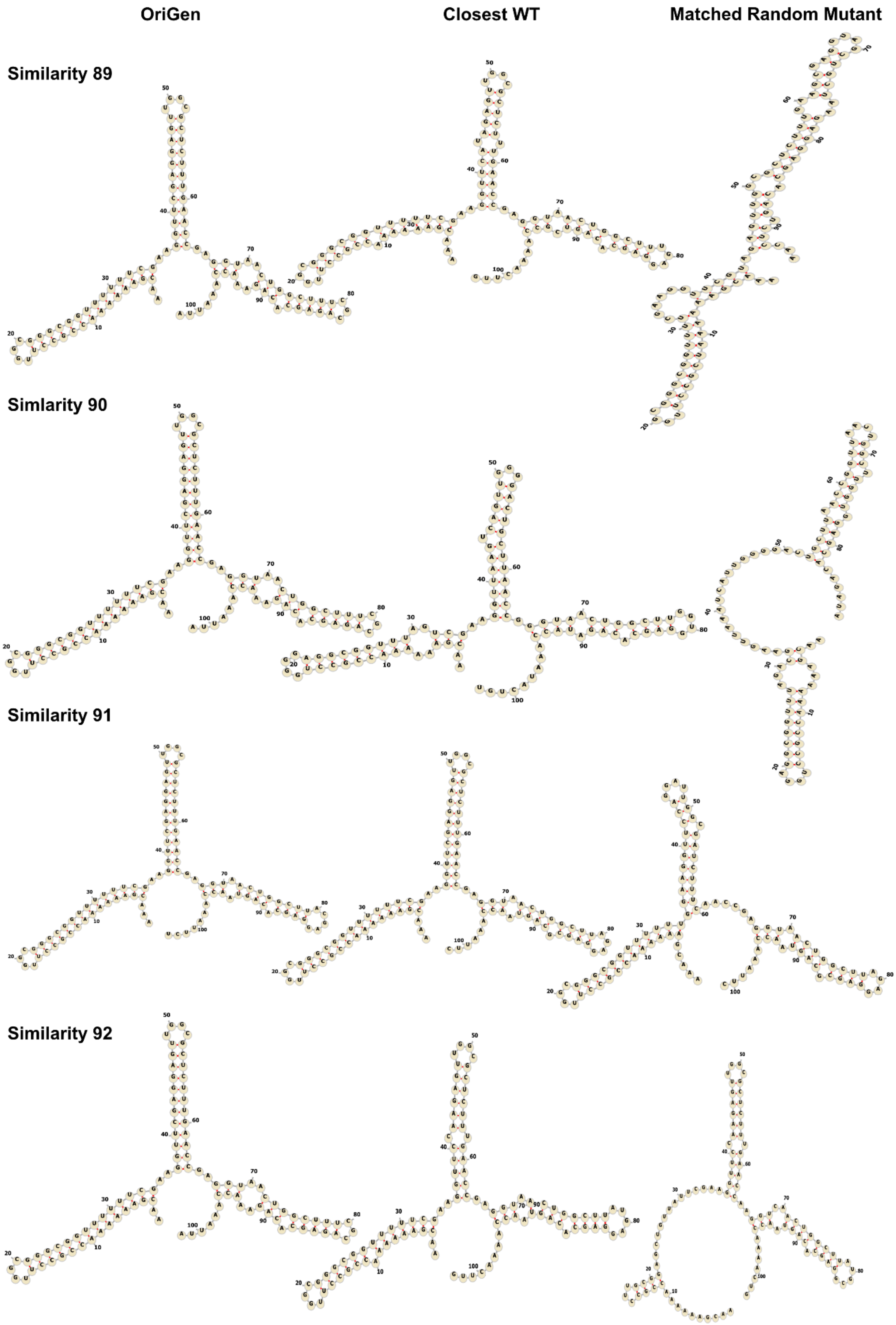
OriGen
Closest WT
Matched Random Mutant
Similarity 89
Simlarity 90
Similarity 91
Similarity 92

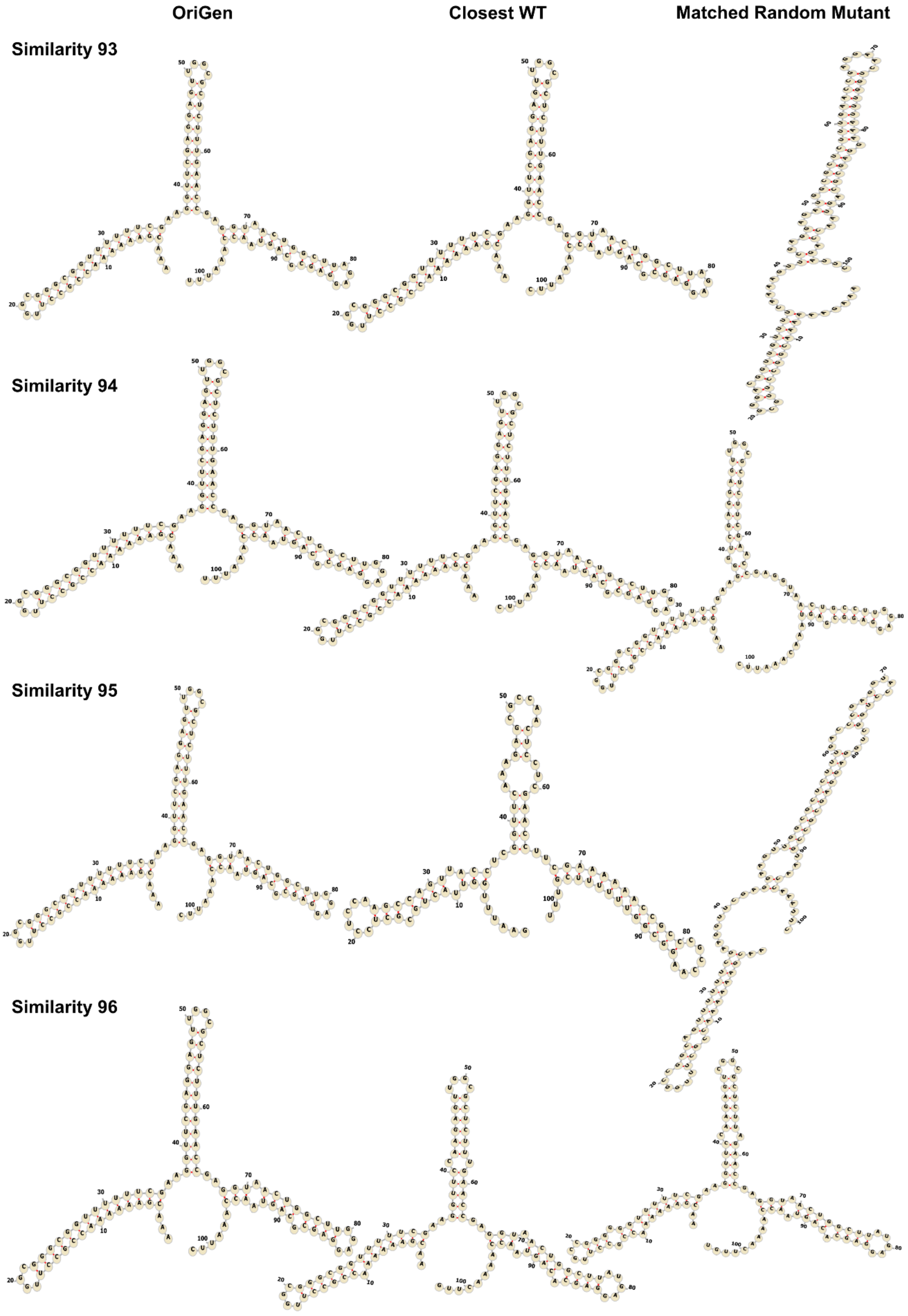
OriGen
Closest WT
Matched Random Mutant
Similarity 93
Similarity 94
Similarity 95
Similarity 96

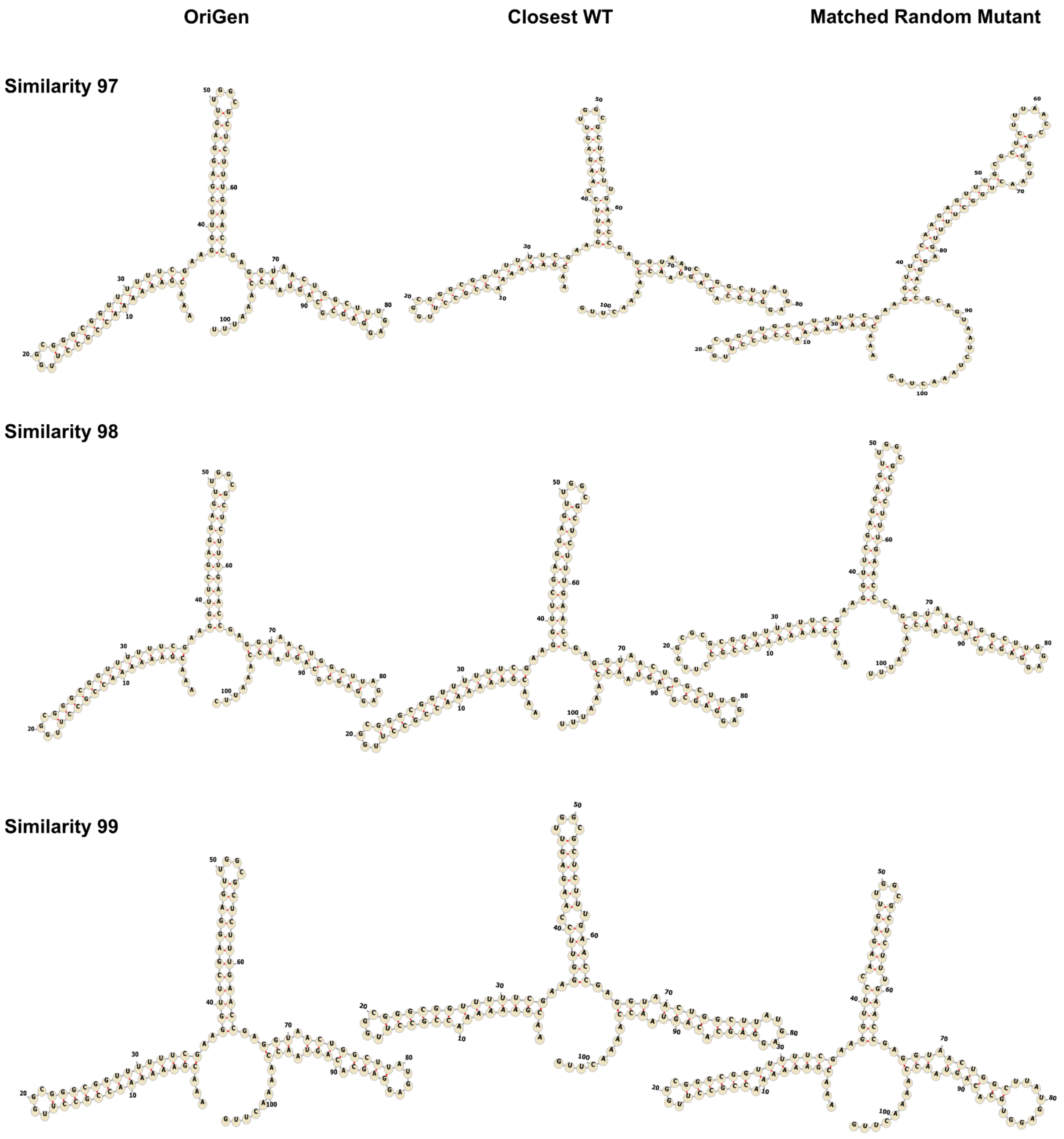


**Supplemental Figure 2:** Predicted structures of ColE1 RNAI for closest wild type (WT) origin, OriGen generated, and matched random mutant controls. Structures are arranged by sequence similarity to the full wild-type origin sequence, as determined using the Needleman-Wunsch algorithm.

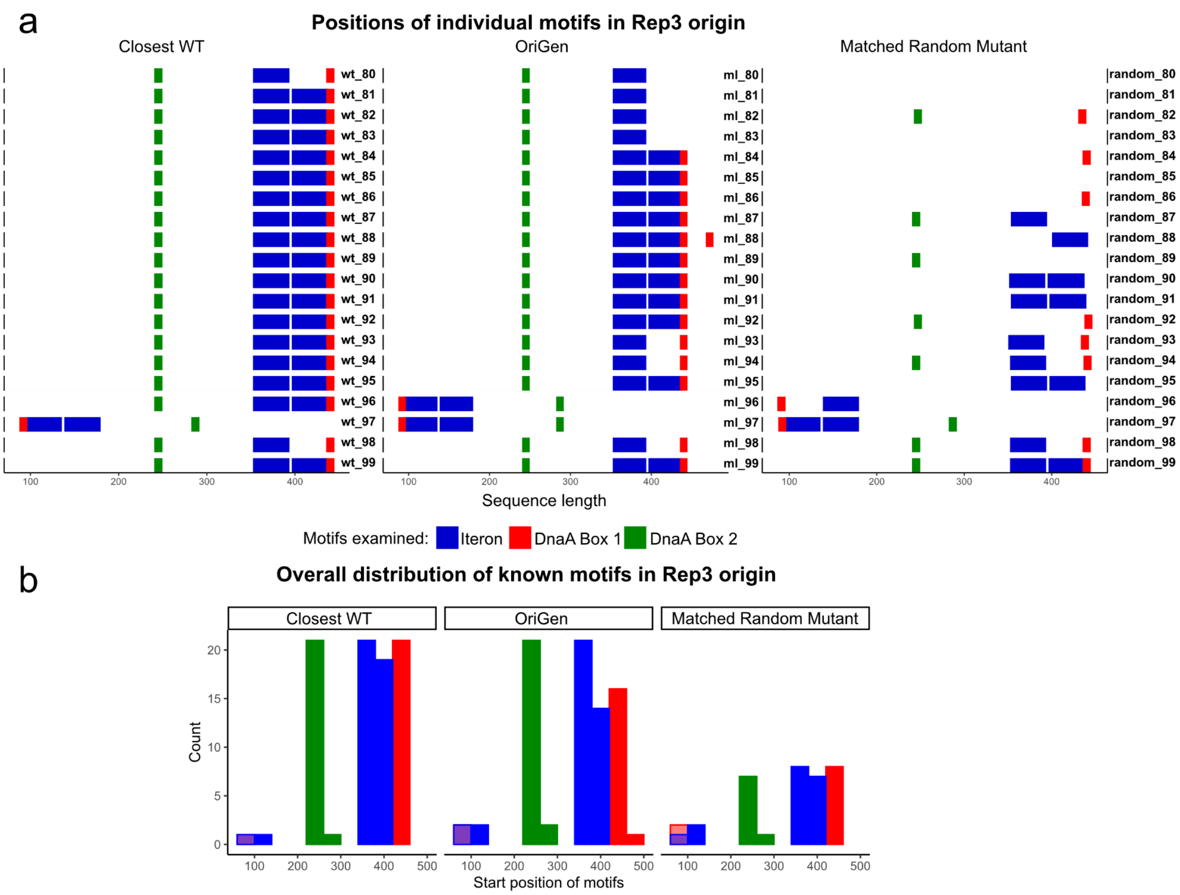


**Supplemental Figure 3: Detection of motifs in Rep3 origins. a,** Known motifs identified in individual origin sequences ordered by Needleman-Wunsch similarity. **b,** Overall counts of motifs identified in the closest wild-type (WT) origin, OriGen generated, and matched random mutant controls. Iteron motifs are colored blue, DnaA box 1 is red, DnaA box 2 is green.

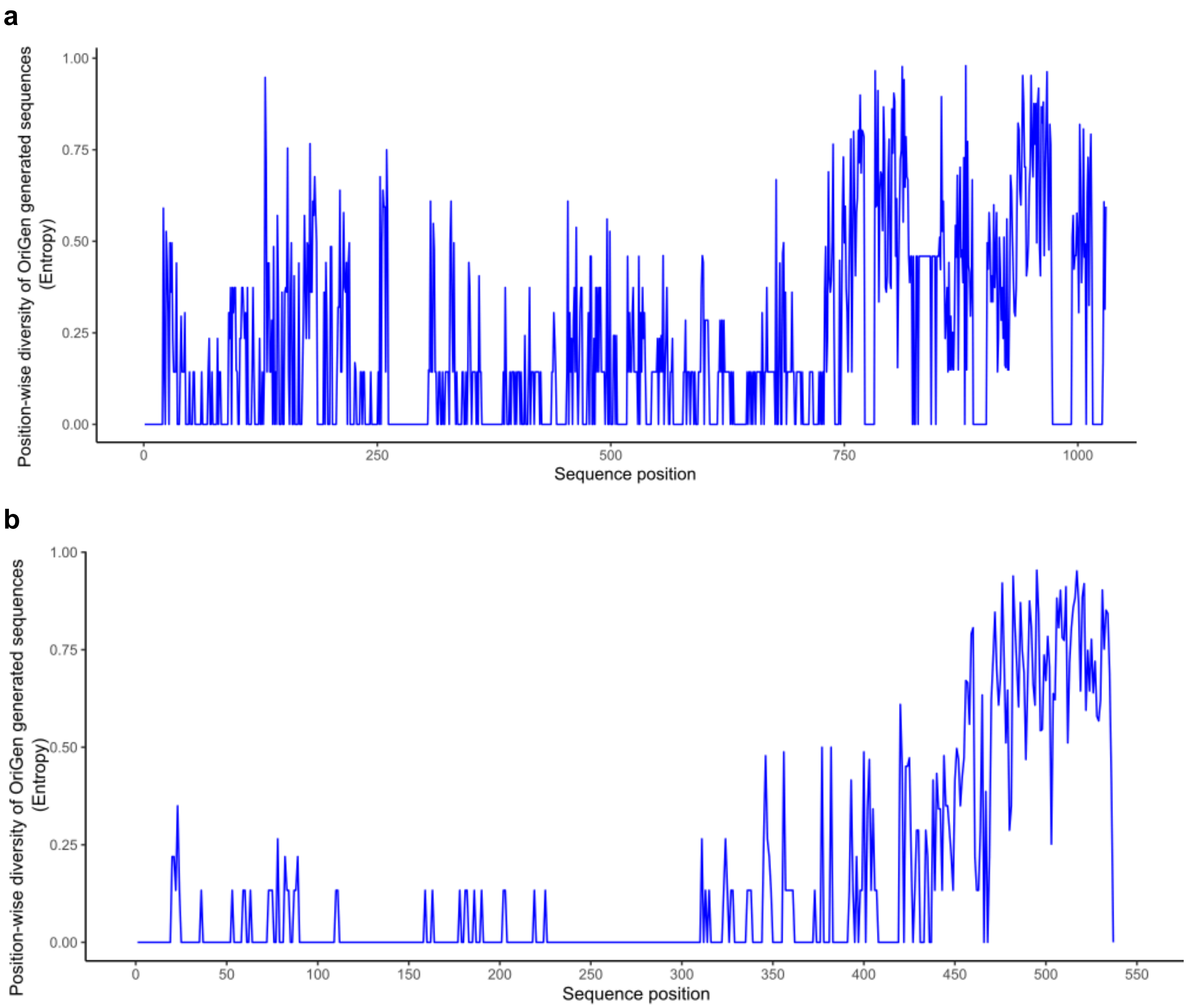


**Supplemental Figure 4: Sequence entropy analysis of OriGen-generated origins. a,** Position-wise Shannon entropy plot of aligned OriGen-generated ColE1-type origins **b,** Corresponding analysis for Rep3-type origins. Lower entropy values indicate higher conservation across sequences.

| Origin type | Similarity bracket | OriGen BLAST pident | OriGen BLAST query coverage | OriGen Needleman % similarity | Random Needleman % similarity |
|---|---|---|---|---|---|
| ColE1 | 80 | 95.19 | 69.71 | 80.28 | 81.42 |
| ColE1 | 81 | 95.39 | 79.95 | 80.83 | 81.88 |
| ColE1 | 82 | 95.41 | 70.11 | 81.88 | 82.97 |
| ColE1 | 83 | 97.37 | 77.76 | 82.93 | 83.31 |
| ColE1 | 84 | 96.84 | 77.76 | 83.8 | 85.19 |
| ColE1 | 85 | 96.62 | 76.81 | 84.82 | 85.24 |
| ColE1 | 86 | 97.31 | 81.17 | 85.6 | 85.73 |
| ColE1 | 87 | 99.12 | 77.76 | 87.08 | 87.02 |
| ColE1 | 88 | 92.07 | 95.53 | 87.64 | 88.27 |
| ColE1 | 89 | 92.24 | 94.41 | 88.61 | 88.71 |
| ColE1 | 90 | 91.16 | 100.61 | 90 | 90.08 |
| ColE1 | 91 | 91.06 | 100.68 | 90.92 | 90.79 |
| ColE1 | 92 | 92.7 | 100.68 | 91.98 | 91.98 |
| ColE1 | 93 | 93.2 | 100.27 | 92.55 | 92.55 |
| ColE1 | 94 | 94.17 | 100.55 | 94.05 | 94.05 |
| ColE1 | 95 | 94.7 | 100.41 | 94.58 | 94.72 |
| ColE1 | 96 | 96.47 | 100.14 | 96.47 | 96.61 |
| ColE1 | 97 | 96.74 | 100.14 | 96.74 | 96.74 |
| ColE1 | 98 | 97.96 | 100 | 97.96 | 97.96 |
| ColE1 | 99 | 98.64 | 100.14 | 98.64 | 98.64 |
| Rep3 | 80 | 91.1 | 90.79 | 80.59 | 81.75 |
| Rep3 | 81 | 100 | 78.72 | 81.91 | 82.12 |
| Rep3 | 82 | 100 | 78.72 | 82.02 | 82.59 |
| Rep3 | 83 | 98.45 | 84.77 | 83.98 | 83.78 |
| Rep3 | 84 | 98.66 | 84.56 | 84.86 | 86.02 |
| Rep3 | 85 | 99.78 | 84.56 | 85.74 | 86.07 |
| Rep3 | 86 | 98.08 | 88.14 | 86.94 | 87.45 |
| Rep3 | 87 | 99.34 | 85.31 | 87.88 | 87.77 |
| Rep3 | 88 | 98.91 | 86.44 | 88.47 | 88.79 |
| Rep3 | 89 | 98.46 | 85.5 | 89.45 | 90.15 |

**Supplemental Table 1: Detailed sequence similarity metrics for experimentally tested origins.** Alignment and BLAST metrics for OriGen-generated sequences and matched random mutant controls. Each row shows the origin type, similarity bracket, BLAST percent identity (pident) and query coverage of the top hit when comparing OriGen-generated sequences against the training dataset, Needleman-Wunsch global alignment similarity score between OriGen-generated sequences and their best matching wild-type sequence identified by BLAST, and Needleman-Wunsch similarity score between random mutant controls and the same wild-type sequence. Random mutants contain identical numbers and types of mutations as their OriGen-generated counterparts but with random positional distribution.
